# Comprehensive variant effect predictions of single nucleotide variants in model organisms

**DOI:** 10.1101/313031

**Authors:** Omar Wagih, Bede Busby, Marco Galardini, Danish Memon, Athanasios Typas, Pedro Beltrao

**Affiliations:** E uropean Molecular Biology Laboratory, European Bioinformatics Institute, Wellcome Trust Genome Campus, Hinxton, CB10 1SD, Cambridge, UK; European Molecular Biology Laboratory, Genome Biology Unit, 69117 Heidelberg, Germany

## Abstract

The effect of single nucleotide variants (SNVs) in coding and non-coding regions is of great interest in genetics. Although many computational methods aim to elucidate the effects of SNVs on cellular mechanisms, it is not straightforward to comprehensively cover different molecular effects. To address this we compiled and benchmarked sequence and structure-based variant effect predictors and we analyzed the impact of nearly all possible amino acid and nucleotide variants in the reference genomes of *H. sapiens*, *S. cerevisiae* and *E. coli*. Studied mechanisms include protein stability, interaction interfaces, post-translational modifications and transcription factor binding sites. We apply this resource to the study of natural and disease coding variants. We also show how variant effects can be aggregated to generate protein complex burden scores that uncover protein complex to phenotype associations based on a set of newly generated growth profiles of 93 sequenced *S. cerevisiae* strains in 43 conditions. This resource is available through mutfunc, a tool by which users can query precomputed predictions by providing amino acid or nucleotide-level variants.

## Introduction

One of the key challenges of biology is to understand how genetic variation drive changes in phenotypes. Genome-wide association studies (GWASs) have made progress in identifying causal genetic loci and over the past decade, a large number of associations have been made between genetic variation and phenotypic traits including disease risk (Welter *et al*, 2014). However, GWASs are typically limited in their ability to explain the underlying mechanism that is influenced by the variant in question. This missing mechanistic layer severely limits our understanding of how variants cause phenotypic variability.

Variants occurring in coding and noncoding regions can influence a diversity of molecular functions. For instance, non-coding variants can affect chromatin accessibility (Kumasaka *et al*, 2016), splice sites (Xiong *et al*, 2015), and epigenetic modifications (Rintisch *et al*, 2014). Coding variants can affect post-translational modification (PTM) sites (Reimand *et al*, 2015; Wagih *et al*, 2015), protein folding and stability (Lorch *et al*, 2000), protein interaction interfaces (Engin *et al*, 2016), sub-cellular localization (Björses *et al*, 2000), and introduce premature stop codons. Understanding the disrupted biological mechanisms underlying genetic variation is key to many applications in genetics such as genetically engineering organisms, assessing drug efficacy and drug discovery (Nelson *et al*, 2016; Labaudinière, 2002; Lutz, 2010).

The ability to predict the degree to which genetic variation would alter such mechanisms offers a time and cost-effective alternative over experimental approaches to prioritize variants of interest and to facilitate the understanding of the mechanisms underlying causal variants. A multitude of *in silico* predictors aimed at predicting such effects have been proposed (Wagih *et al*, 2015; Schymkowitz *et al*, 2005; Adzhubei *et al*, 2010; Kumar *et al*, 2009), yet they often require significant computational power, expertize and time to be used. Furthermore, each of the currently available tools do not comprehensively provide predicted effects across different molecular mechanisms (i.e disruption of stability, interfaces, TF binding etc).

Accordingly, we have compiled and benchmarked commonly-used sequence and structure-based predictors of mutational consequences and predicted the effect of nearly all possible variants in the reference genomes of *H. sapiens*, *S. cerevisiae*, and *E. coli*. The impact of variants was measured in the context of conserved protein regions, protein stability, protein-protein interaction (PPI) interfaces, PTMs, kinase-substrate interactions, short linear motifs (SLiMs), start and stop codons, and transcription factor (TF) binding sites (TFBSs). This resource is available through the mutfunc resource (http://mutfunc.com/), which allows for prioritization of variants while providing insight into the altered mechanisms.

To demonstrate the utility of mutfunc, we assessed variants of uncertain clinical significance (VUSs) in *H. sapiens*. We further applied mutfunc to publically available variants for yeast *S. cerevisiae* strains to generate protein complex burden scores. We then phenotyped 93 sequenced *S. cerevisiae* strains in 43 conditions and utilized burden scores to associate protein complexes to phenotypes. This yielded associations that would not be possible through traditional variant-based GWAS approaches. Mutfunc is a computational resource that will facilitate the study of the mechanistic impacts of genetic variation.

## Results

### Functional genomic regions display evolutionary constraint across yeast and human individuals

In order to set up the variant effect prediction approaches we first derived, for *E. coli*, *S. cerevisiae* and *H. sapiens*, molecular information such as: experimental and homology based protein structural models for individual proteins and protein interfaces, TF binding sites, protein kinase targets sites, post-translational modification sites and linear motif regions (**Methods**). Structural models were used to identify interface residues and residues with different surface accessibility. Given that functionally-relevant regions of the genome are are under evolutionary constraint we took the opportunity to use this large collection of functional regions to test if these tend to be depleted of natural variants. For yeast 896,772 natural variants and their allele frequencies were compiled from 405 yeast strains (Strope *et al*, 2015; Zhu *et al*, 2016; Gallone *et al*, 2016; Bergström *et al*, 2014), of which 478,857 were coding variants. For human, over 3.2M coding variants from over 65,000 individuals were obtained from the ExAC consortium (Lek *et al*, 2016).

Natural variants were mapped to 9,837 protein structures and homology models (n=6,737 human, n=3,100 yeast) and the residues were binned according to relative surface accessibility (RSA). Similarly, 9,883 structures (n=7,693 human, n=2,190 yeast) for protein interaction pairs were obtained from Interactome3D and the difference in surface accessibility (∆RSA) between the unbound and bound complex were determined to identify interface residues, corresponding to those with the highest ∆RSA (**Methods**). The number of variants per position of each bin of RSA and ∆RSA were compared to counts observed in random positions in the protein, permuted 1,000 times. Fewer variants were found in buried regions and interface regions when compared to exposed regions in both yeast and human (**Figure 1a** p<1.28x10^−34^ and **1b** p<2.28x10^−33^). To study variation at 296,147 and 26,560 human and yeast PTM sites the variant counts over random expectation was calculated for a window of +/-5 residues flanking the PTM positions. The level of constraint was different across PTM types (**Figure 1c**) with ubiquitin showing the lowest level of constraint. Interestingly, the level of constraint for PTMs increases with the number of other neighboring PTMs present in a 10 amino acid window (**Figure 1d**) suggesting that the clustering of PTMs may have important biological functions such as cross-talk regulation (Beltrao *et al*, 2013).

**Figure 1.**
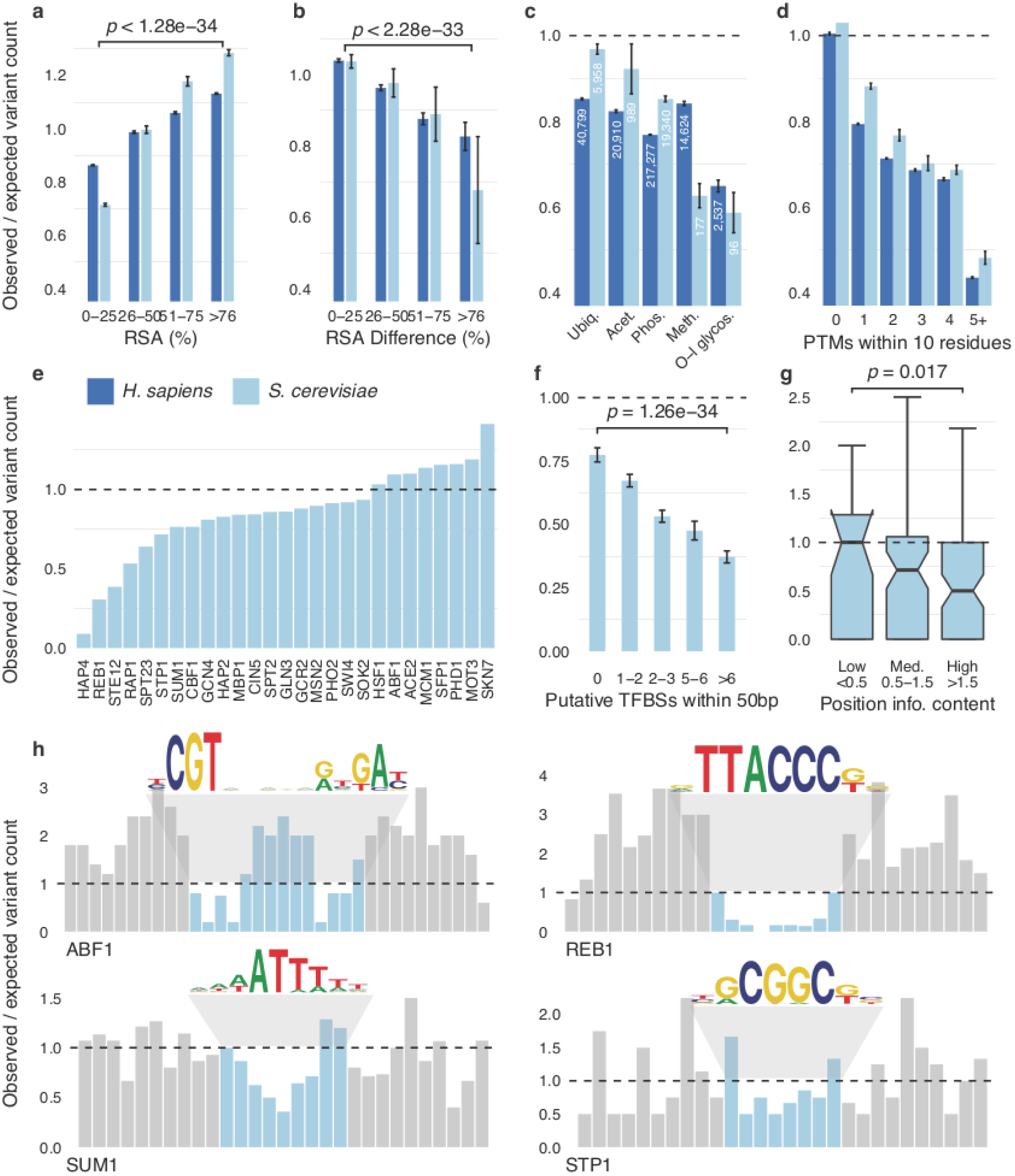
Population level sequence constraint in genome functional elements. The level of sequence constraint was estimated using a ratio of the counts of genome variants across individuals of yeast and human compared with a random control regions for different functional elements (a) Regions buried within a protein structure with a low RSA typically exhibit higher evolutionary constraint. Similarly, (b) regions buried within interaction interfaces exhibit a high ∆RSA and demonstrate stronger sequence constraints. P-values represent a one-sided Wilcoxon test. (c) sequence constraint on PTMs, where numbers reflect the number of PTM sites for each modification. (d) PTMs with a higher number of neighbouring PTMs are show stronger constraint. (e) Variability in constraint amongst bindings sites for TFs with at least 40 sites. (f) TFBSs that coexist with other binding sites are under stronger constraint. P-value shown is computed using a one-sided Wilcoxon test (g) Position-specific constraint shows that positions of higher relevance for binding in TFs with at least 20 sites are under stronger constraint. P-value shown is computed using a one-sided Kolmogorov-Smirnov test. (h) four examples where the bar plots reflect the position-specific constraint in (blue) and around (grey) the binding site, along with sequence logos for the binding specificities.

We next analysed non-coding variation at putative TF binding sites for *S. cerevisiae* that were predicted using a combination of TF specificity models, TF knockout gene expression studies and TF ChIP-seq or ChIP-chip data (**Methods**). A total of 4,523 potential binding sites were identified across 93 TFs of *S. cerevisiae*. We computed the ratio between the variant counts within the predicted binding sites to that of random genomic sites of the same length and within the same ChIP regions. By combining the analysis across all putative binding sites of each TF we observed that binding sites for some TFs are generally more constrained than others (**Figure 1e**). Those with higher levels of constraint include HAP4, a global regulator of respiratory genes and general transcriptional regulators such as REB1 and RAP1. At the level of individual TF binding sites we observed that those found within clusters of binding sites tended to show higher levels of constraints than isolated sites (**Figure 1f**). Additionally, the TF binding positions for each TF were stratified according to their importance for binding as measured by the position-specific information content (IC) of the TF specificity position weight matrices. In accordance with expectation, positions with high IC, that correspond to positions that are important for binding, tend to have fewer variants than less important positions (**Figure 1g**). Position-specific constraint for individual TFs highlights this difference between high and low IC positions (**Figure 1h**).

Overall these results provide an overview of how population level variation differs across diverse set of genome functional elements and recapitulates findings from analysis of specific types of functional elements (Spivakov *et al*, 2012; de Beer *et al*, 2013; Reimand *et al*, 2015). Additionally, it suggests that our collection of functional elements (e.g. structures, interfaces, PTMs and TF binding sites) shows evolutionary constraints and therefore can be used further for the establishment of the variant effect prediction pipeline.

### A comprehensive resource of mechanistic effects of single nucleotide variants

We sought to better understand the mechanistic impact of point mutations affecting the above described functional elements. To do this, a set of commonly-used predictors were used to assess the impact of every possible single amino acid or nucleotide substitution across *H. sapiens*, *S. cerevisiae*, and *E. coli*, where applicable. We performed a large scale computational estimation of the impact of variants on conserved protein regions, protein stability, protein interaction interfaces, kinase-substrate phosphorylation and other PTMs, linear motifs, TFBSs and start and stop codons (illustrated in **Figure 2a**, **Methods**). These results were deposited in the mutfunc resource, which offers a quick and interactive way by which users can gain predicted mechanistic insight for variants of interest.

**Figure 2.**
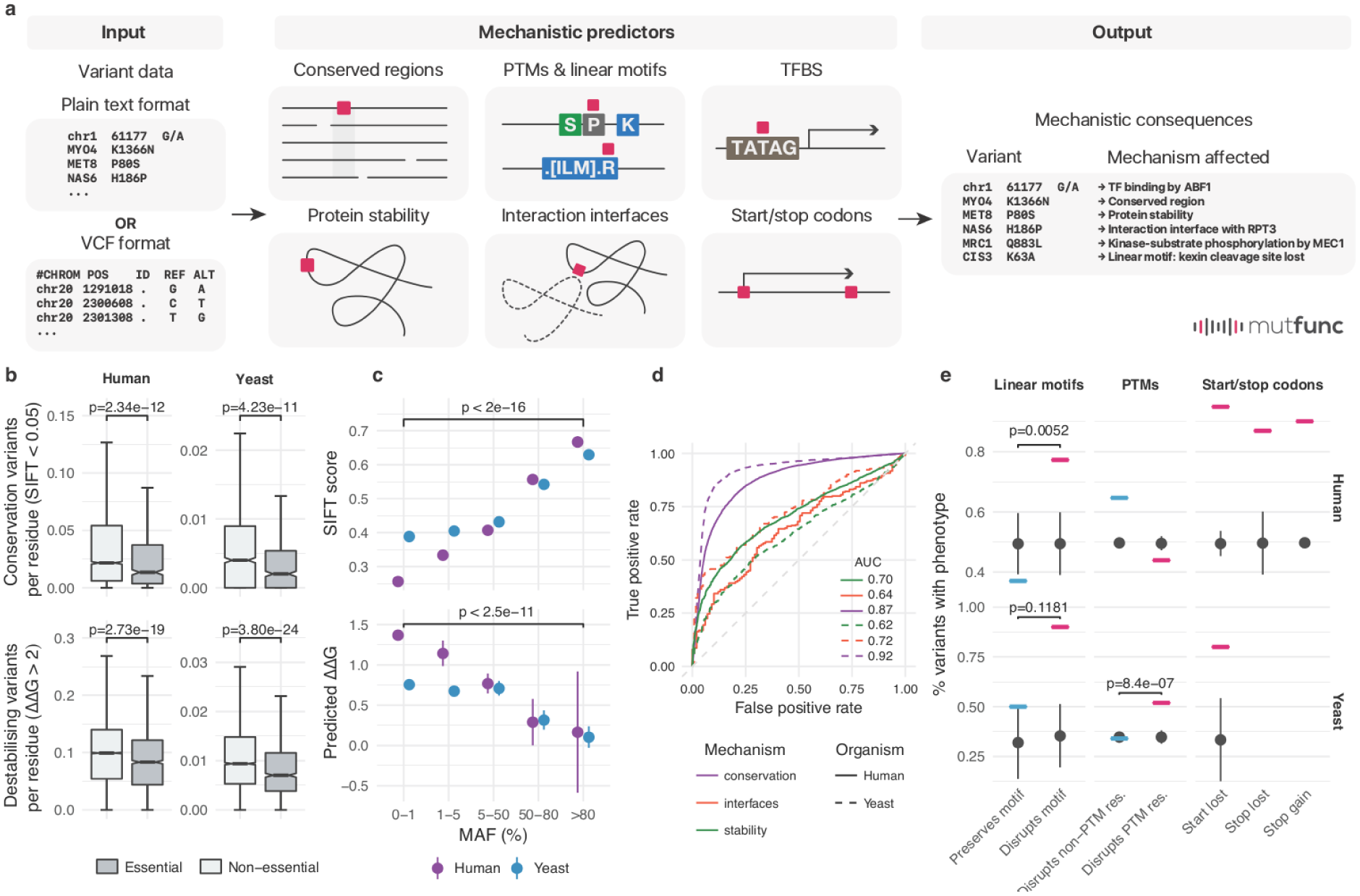
The mutfunc resource and benchmarking of underlying variant effect predictors. a) The mutfunc interface provides an intuitive, user-friendly way by which users can query the resource using DNA or protein substitutions provided in plain text format or the variant call format (VCF). The impact of variants across different mechanisms are provided with information on impact strength in downloadable format and/or protein structural views. (b) The fraction of variants predicted to affect a conserved of structural important residues for essential and non-essential genes. (c) mean SIFT scores and predicted ∆∆G values for human and yeast variants within different MAF bins. Error bars represent the standard error and p-values are calculated based on a one-sided Wilcoxon test. (d) Benchmarking of the capacity of different predictors to discriminate between pathogenic and benign variants. (e) the proportion of pathogenic versus benign variants that disrupt or not different functional annotations (SLiMs, PTMs or stop gains/losses).

To measure the impact on conserved regions, we constructed 29,027 multiple sequence alignments for proteins of the three organisms (n=19,497 *H. sapiens*, n=5,498 *S. cerevisiae*, n=4,032 *E. coli*), and used the SIFT algorithm (Ng & Henikoff, 2003) to assess the impact of all possible 291.7M protein coding variants (n=212.2M *H. sapiens*, n=53.4M yeast, n=26.1M *E. coli*). To measure the impact on protein stability, the FoldX algorithm (Schymkowitz *et al*, 2005) was applied to 17,893 structures (including homology models) across the three organisms, and precomputed effects of 66.3 million protein coding substitutions (n=42.7M *H. sapiens*, n=10.3M *S. cerevisiae*, n=13.4M *E. coli*, **Methods**). We identified interface residues in 10,675 structures of binary PPIs from Interactome3D across the three organisms and similarly applied FoldX to compute the effects of 11.2M possible interface mutations on binding stability (n=7.2M *H. sapiens*, n=2.3M *S. cerevisiae*, n=1.6M *E. coli*). To identify variants that could impact kinase-substrate sites, we used MIMP (Wagih *et al*, 2015) to predict the impact of all possible 541,161 variants (n=485,736 *H. sapiens*, n=55,425 *S. cerevisiae*) falling within ±5 residues of a known kinase-substrate phosphorylation site (phosphosite) on a kinase’s specificity. Specificities for 56 kinases in *H. sapiens* and 46 kinases in *S. cerevisiae* were considered. Kinase-phosphosite relationships for *E. coli* are not well established and cannot be scored in the same way. For all other PTMs such as methylation, ubiquitination, and acetylation for which we do not have explicit flanking sequence specificity models, a variant was considered damaging if it directly altered the modified site. This resulted in a total of 6.3M possible variants that could alter such PTM sites across the three organisms (n=5.8M *H. sapiens*, n=537,434 *S. cerevisiae*, n=9,177 *E. coli*). For linear motif information, not available for *E. coli*, we gathered 1,668 experimentally identified linear motifs (n=1,525 *H. sapiens*, n=143 *S. cerevisiae*), along with their derived regular expression pattern from the ELM database (Dinkel *et al*, 2012) and computed the impact of all possible 226,920 variants (n=205,120 *H. sapiens*, n=21,800 *S. cerevisiae*) on binding patterns. Finally, for TFBSs, for organisms without well-defined functional TFBSs (*H. sapiens* and *S. cerevisiae*), we defined putative TF-gene regulatory network using TF-knockdown expression data and/or ChIP-seq/ChIP-chip (**Methods**). We then used PWMs to identify putative binding sites, and predict the impact (**Methods**) of all possible 3.6M variant substitutions (n=3.3M *H. sapiens*, n=236,382 yeast, n=46,768 *E. coli*) on specificities of 217 TFs (n=72 *H. sapiens*, n=104 *S. cerevisiae*, n=41 *E. coli*).

These pre-computed variant effect predictions constitute a resource that can be used in diverse ways. In the next sections we benchmark this resource and illustrate some of its possible applications.

### Functionally important positions are enriched in predicted deleterious variants

In order to benchmark the variant effect predictions that underlie the mutfunc resource we first asked if essential genes would harbour fewer natural variants that are predicted to be deleterious. Essential genes in yeast (Giaever & Nislow, 2014) and human (Blomen *et al*, 2015) consistently demonstrated significantly lower frequencies of variants predicted to affect conserved sites (sift score < 0.05, p=1.04x10^−46^ human, p=1.52x10^−22^ yeast, **Figure 2b**) and protein stability (∆∆G pred > 2, p=1.82x10^−12^ human, p=7.07x10^−10^ yeast, **Figure 2b**). Variants of higher allele frequency in the population are expected to be less impactful and in accordance to this we observed an increase in deleterious scores, as predicted from SIFT and FoldX, for variants of lower allele frequencies (**Figure 2c**). In addition to allele frequencies we analyzed mutations that are known to be deleterious. For *H. sapiens* we used 34,600 variants annotated to be pathogenic (n=17,167) or benign (n=17,433) from the ClinVar (Landrum *et al*, 2014). For *S. cerevisiae* we used 8,083 variants consolidate by Jelier et al. (Jelier *et al*, 2011)as either tolerated (n=5,271) or affecting function (n=2,812) (**Methods**). The different predictors consistently discriminated tolerated from pathogenic variants as measured by the area under the receiver operating characteristic curve (AUC). SIFT performed the best at discriminating pathogenic variants from benign (AUC *H. sapiens* = 0.87, *S. cerevisiae* = 0.92), followed by FoldX interfaces (AUC *H. sapiens* = 0.64, *S. cerevisiae* = 0.72) and FoldX stability (AUC *H. sapiens* = 0.70, *S. cerevisiae* = 0.62, **Figure 2d**).

For other heuristic-based predictors such as SLiMs, PTMs or stop gains/losses we compared the proportion of pathogenic versus benign variants that disrupt or not the annotation. Despite the low number of pathogenic variants overlapping with these features we observed an enrichment of pathogenic variants for mutations that disrupt such features (**Figure 2e**). The only exceptions were for PTM disrupting variants in human and for linear motif disrupting variants in yeast. In contrast, there were significant differences for the enrichment of pathogenic variants disrupting human linear motifs (p=5.23x10^−3^) and yeast PTM sites (p=8.44x10^−7^). For some of annotations the lack of statistical significance may be due to the small number of testable variants.

The results here demonstrate that the predictors used in mutfunc are generally capable of enriching for variants of functional significance. The resource can be used to prioritize variants according to the degree of pathogenicity as well as provide molecular mechanisms affected.

### Predicting mechanistic impacts of variants of uncertain significance

Variants that have been identified through disease related genetic testing but are yet to be deemed benign or pathogenic are termed variants of uncertain significance (VUS). The interpretation of such variants is a common challenge in genetics, one that is often aided by computational predictors. A total of 64,692 variants labelled with “uncertain significance” were collected from ClinVar (Landrum *et al*, 2014). VUSs were annotated using mutfunc and 21,584 variants were predicted impactful by at least one of the mechanistic predictors, not including SIFT (n=7,547 stability, n=751 interfaces, n=139 linear motifs, 2,372 PTMs, 57 kinase-binding). From these we focused on variants predicted to impact the structural integrity of proteins (stability and interaction interfaces) since they hold the highest coverage. Of the VUSs predicted to interfere with interface or protein stability we retained those in which (1) the protein also harbours a known pathogenic variant with the same predicted structural impact and (2) both the pathogenic variant and VUSs are identified in patients carrying the same disease. This allows us to connect VUSs to pathogenic variants via shared altered mechanisms and affected phenotypes. We demonstrate a few examples of VUSs that are predicted to alter binding (**Figure 3a, 3b and 3c**) or structural stability (**Figure 3d and 3e**). For instance, primary hyperoxaluria is a disease caused primarily by mutations in GRHPR, a glyoxylate and hydroxypyruvate reductase (Cramer *et al*, 1999; Cregeen *et al*, 2003) and its enzymatic activity requires homodimerization (Booth *et al*, 2006). For this enzyme the variants R302H and E113K have been implicated in primary hyperoxaluria, are known to be pathogenic and are predicted to impact on binding stability (**Figure 3a**, ΔΔG > 2.15). We can reason that other variants in patients of the same disease impacting on GRHPR homodimerization should be equally pathogenic. For example, the variant R171H is predicted to impact a conserved region as well as the homodimerization stability (**Figure 3a**, ΔΔG = 2.19, *s* < 0.018) and found in primary hyperoxaluria patients. Although R171H is of uncertain significance our analysis strongly suggests that it is pathogenic through the same molecular mechanism as R302H and E113K. Similarly compelling examples are found for other proteins such as fumarate hydratase (**Figure 3b**) and lamin (**Figure 3c**).

**Figure 3.**
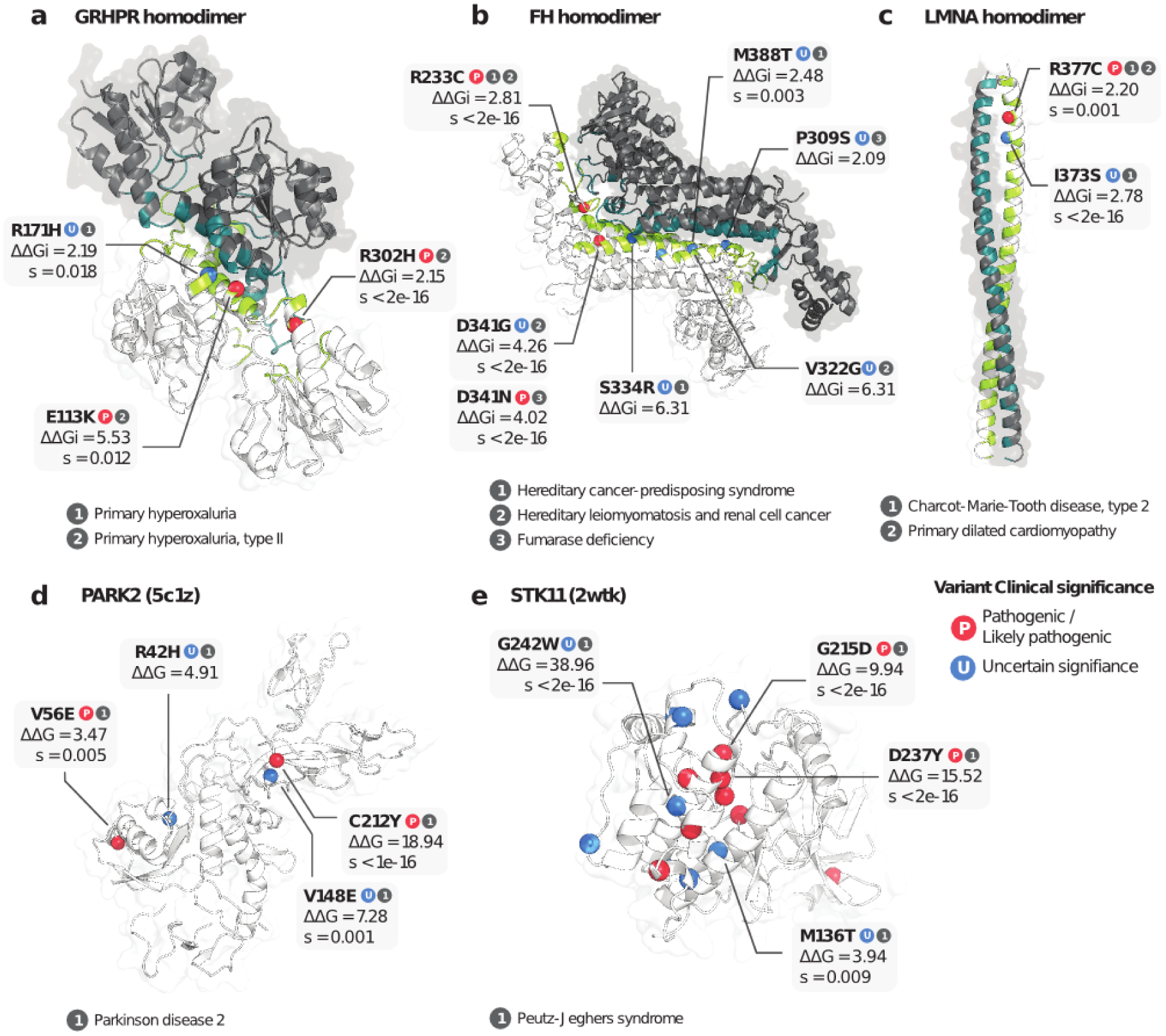
Analysis of variants of uncertain clinical significance using mutfunc. (a-c) Three examples of interaction interfaces containing variants predicted to impact binding stability. Subunits of the interaction complex are coloured in dark grey and white, and respective interface residues in dark green and green. (b) Two examples of variants predicted to impact protein stability. Pathogenic variants are labelled “P” in red, and VUSs “U” in blue.

Similar to interface variants, we analysed variants that destabilise the protein structure. We identified 1,182 VUSs predicted to alter stability in proteins containing pathogenic variants also predicted to be destabilizing. For instance, the ubiquitin ligase PARK2, implicated in Parkinson’s disease, contains pathogenic variants predicted to impact on its stability. For this protein two rare VUSs (R42H, V148E) identified in Parkinson’s disease patients are similarly predicted to destabilise the protein (ΔΔG > 4.7, **Figure 3d**) and are therefore likely to be pathogenic. In the tumour suppressor serine/threonine-protein kinase STK11, pathogenic and VUS identified in Peutz-Jeghers syndrome patients can be similarly linked (**Figure 3e**).

The analysis here demonstrates how mutfunc could be applied to systematically prioritize pathogenic variants through altered mechanisms that may be the molecular cause of the phenotype.

### *S cerevisiae* strain genomic differences are a poor predictor of phenotypic similarity

We sought to illustrate the use of mutfunc for genotype-to-phenotype association analysis. Using *S. cerevisiae* as a case study, we first phenotyped growth for a panel of 166 strains in 43 conditions (**Methods**). Colony sizes for strains were quantified, normalised and scored relative to all strains in a condition to produce a phenotypic measure defined as the S-score (Collins *et al*, 2006). Positive and negative values indicate higher or lower than expected growth for a given strain and a specific condition (**Methods**). S-scores for biological replicates demonstrated a high degree of concordance (*r* = 0.91, *p* < 2.22x10^−16^, **Figure 4a**) suggesting a high degree of confidence in phenotypic measurements. S-scores for each strain and growth condition are provided in **Supplementary Table 1**.

**Figure 4.**
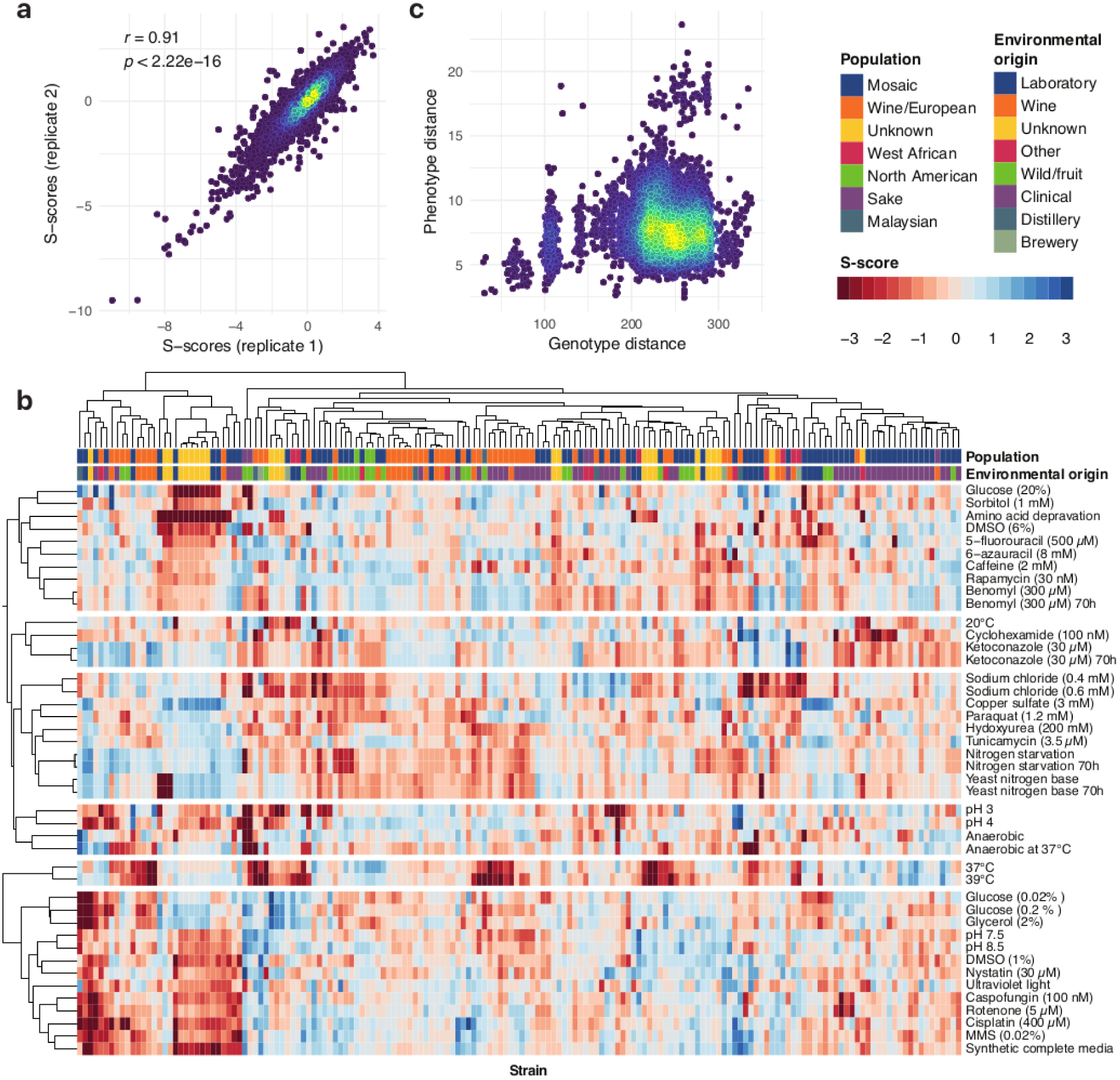
Phenotypic screening of 166 yeast strains. (a) Concordance between replicate s-score measurements. (b) Heatmap of s-scores showing hierarchical clustering of both strains and conditions reveals clusters of phenotypically similar strains and conditions. (c) Comparison of pairwise genotype and phenotype distances between 93 sequenced strains shows little observable correlation.

Hierarchical clustering of growth phenotypes revealed known clusters of related stressors (**Figure 4b**). Clusters of similar phenotypic profiles included for example: UV light, cisplatin and MMS, which are all DNA damaging agents (mean pearson’s *r* = 0.51); nystatin and caspofungin, which interfere with the cell membrane (*r* = 0.49); and caffeine and rapamycin, both involved in TOR signalling (*r* = 0.41). Furthermore, strains belonging to the same population structure (Strope *et al*, 2015) or environmental origin often showed similar phenotypic profiles (**Figure 4b**). Genome sequences were available for 93 of the 166 profiled strains and used to calculate pairwise genomic similarity. As expected, genetic similarity alone is a poor predictor of phenotypic response similarity (**Figure 4c**). This is not unexpected since most genetic variation is expected to be neutral and distantly related strains accumulate variation that may not have an impact on the phenotypes tested.

### Gene and complex disruption scores for genotype-to-phenotype associations

Given that most variants are expected to be neutral we used mutfunc to interpret the observed variants in each strain at the gene-level by computing a total gene burden or disruption score using the mechanistic predictions for conservation (SIFT), protein stability (FoldX) and protein truncating variants (PTVs, including start loss, nonstop and nonsense variants) (**Figure 5a**). Scores produced by predictors are standardised to reflect the likelihood they are deleterious (**Figure 5b**, **Methods**). This allows for effects of rare variants to be combined across different protein positions and predictors into a single probability that the gene is affected (P_AF_ score or burden score) (Jelier *et al*, 2011; Galardini *et al*, 2017).

**Figure 5.**
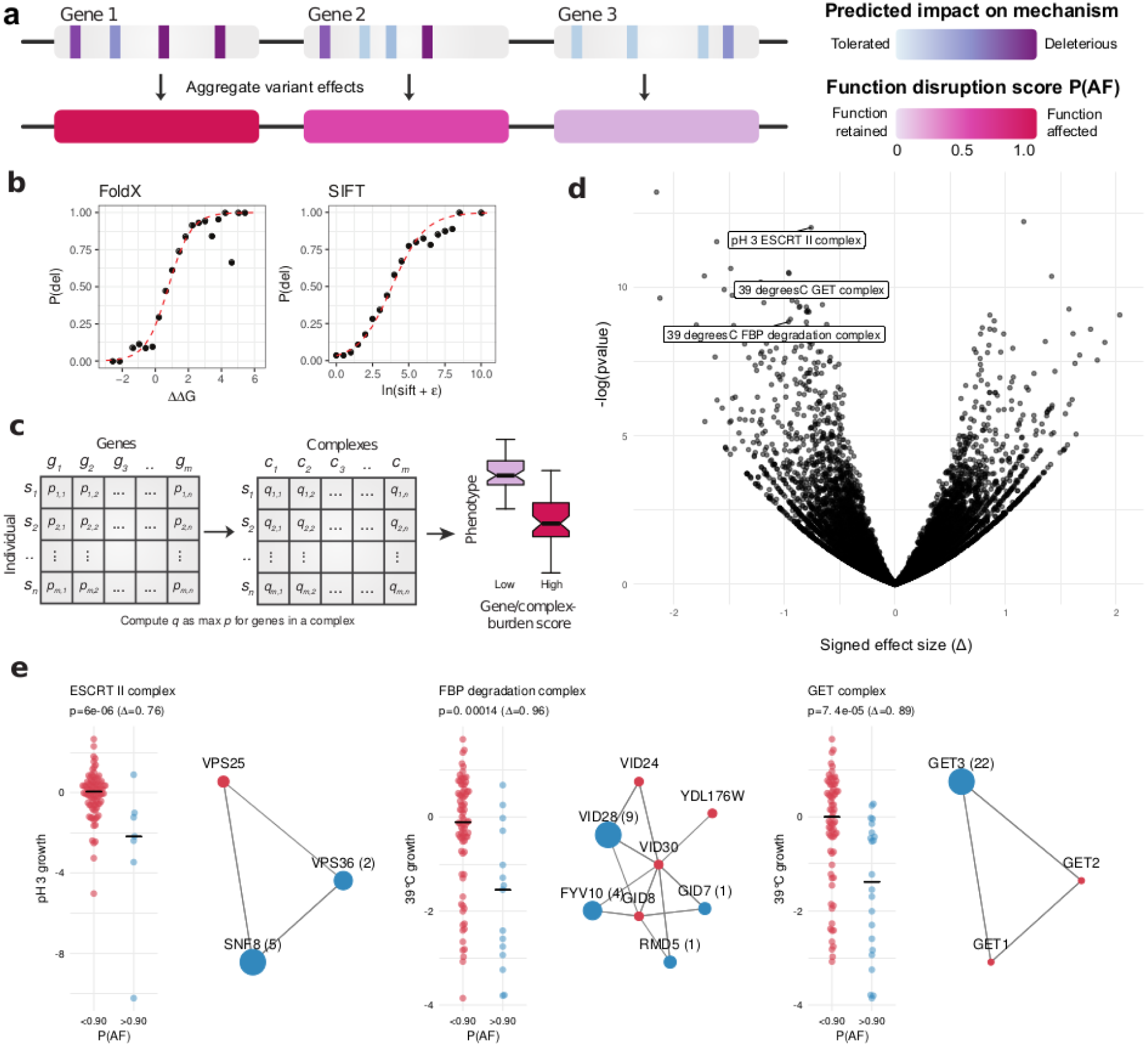
Gene and protein complex level aggregation of variant effects for phenotype association analysis. (a) Diagram demonstrating the aggregation of variant impact. Each variant is first assigned a probability of deleteriousness, which are aggregated at the gene level using the maximum impact. (b) The probability of deleteriousness for FoldX and SIFT was computed by assessing the proportion of deleterious variants in gold-standard data for FoldX and SIFT. A logistic regression model (red line) is fit to compute subsequent probabilities. Protein complex level burden scores was taken to be the maximal burden for any complex member (c) Gene and complex burden scores for each strain, gene/complex-phenotype associations were carried out.

Using the gene level disruption scores we performed phenotype association analysis. Scores were binned based on high (P_AF_> 0.90) or low (P_AF_< 0.90) burden (**Figure 5c**). Associations were carried out for 1,446 genes (with at least three strains containing a P_AF_> 0.90) against growth phenotypes across 43 conditions (**Methods**). All reported p-values were corrected using the false discovery rate (FDR) method and the effect size was computed using the Glass’ Δ approach (**Methods**). We identified 626 statistically significant gene-phenotype associations at p<1x10^−3^ and FDR<10%, with 83% (520/626) being negative i.e. decreased growth (**Supplementary Figure 1a**). Under the assumption that gene function is conserved across strains of *S. cerevisiae* we expected these associations to be enriched in genes that cause a condition specific phenotype when knocked-out. Such association between gene KOs and condition-specific growth phenotypes exist for the lab strain as part of extensive published chemical-genetic studies. We found KO chemical genetic data for 35 of the 43 conditions tested. Of the significant negative associations, only 9% (48/520) are validated by the chemical genetic data. The validation rate increases for higher effect sizes (**Supplementary Figure 1b**) to 13% (38/282) and 24% (15/64) for at Δ > 1 and at Δ > 1.7, respectively. However, based on permutation testing only the enrichment found at large effect sizes (|Δ| > 1.7) was significant (p= 0.03, **Supplementary Figure 1b**).

We next reasoned that protein complex members often act as coherent functional units and that the dysfunction of complex subunits often elicits similar phenotypic outcomes (Collins *et al*, 2007). Therefore, we aggregated the gene-level scores to identify complexes that were potentially defective in a given strain. We performed protein complex level associations focusing on 263 complexes with at least two high burden genes across strains (**Figure 5d**). A total of 122 significant complex-phenotype associations were identified (p<1x10^−3^, FDR < 10%), 48 (39%) of which had a high effect size (∆ > 1). The 48 associations involved 21 conditions and were preferentially negative associations (33 of 48, 69%). We found a significant enrichment in KO chemical genetics data (**Supplementary Figure 1c**) that was significantly higher than observed based on random permutation testing (**Supplementary Figure 1c,** p=0.04). This enrichment is observed only for stringent cut-offs for defining gene-deletion phenotypes from the KO chemical genetic studies (Hillenmeyer *et al*, 2008).

Some examples illustrate how the analysis at the protein complex level may increase power for the identification of associations (**Figure 5e**). For example we found associations between the ESCRT II complex and growth in low pH and the FBP complex, responsible for protein degradation, and high heat (**Figure 5e**). In both cases the individual complex members with high burden scores were not detected by gene-level burden associations, likely due to insufficient recurrency of mutation at the gene-level but in both cases the associations are validated by KO studies. The GET complex represents an example where the association with growth under heat is found both at the gene level and protein complex level due to recurrence of destabilizing mutations in the GET3 gene (**Figure 5e**).

This association analysis indicates that there is value in combining effects of rare variants at the protein and protein complex level to perform association studies. Although the current study is limited due to the relative small number of strains studied, it illustrates how mutfunc can be applied to the study of diverse set of problems.

## Discussion

The mutfunc resource makes use of variant predictors to precompute millions of variant effects across the reference genomes of *H. sapiens*, *S. cerevisiae*, and *E. coli*. These predictors and their performance have been previously described but the large computational effort and the accompanying web-service (mutfunc.com) constitute a resource that facilitates their use. Within mutfunc, conservation effects hold the highest coverage, (*H. sapiens* 98.6%, *S. cerevisiae* 87.9%, and 96.1% *E. coli*) followed by stability (*H. sapiens* 18.9%, *S. cerevisiae* 16.9%, and 49.2% *E. coli*) and interfaces (*H. sapiens* 2.20%, *S. cerevisiae* 2.84%, and 4.45% *E. coli*). Other mechanisms like PTMs and TFBSs have lower coverage. As additional data become available, mutfunc will be updated to improve coverage and future work could expand the set of mechanisms studied such as drug or small-molecule binding sites, RNA-binding interfaces, among others. The effects of variants on molecular and cellular phenotypes is increasingly being probed directly by large-scale mutagenesis experiments (Fowler & Fields, 2014; Weile *et al*, 2017), which will likely result in improved variant effect prediction algorithms (Gray *et al*, 2018). The curation of such experimentally determined effects and the improved algorithms can be integrated in future iterations of mutfunc.

A strength of mutfunc lies in its large set of precomputed SNV effects allowing for genome-wide variants to be rapidly queried. However, within such a framework, combinatorial and potential epistatic effects cannot be precomputed due to a large number of possible combinations. Similarly, many other types of genetic variation such as copy number variations and indels (Chuzhanova *et al*, 2003; Beroukhim *et al*, 2010) have not be considered in mutfunc due to their complex structure. Lastly, many organisms in which genetic variation is commonly studied are not included in mutfunc. These include *M. musculus*, *D. melanogaster* and *A. thaliana*, which contain an abundance of data and could be added in the future.

Understanding how disrupted cellular mechanisms propagate to changes in phenotypes is critical for variant interpretation. We show here how different variants can be integrated using effect predictors and protein complex annotations to perform genotype-to-phenotype associations for full genome sequences. In addition, we and others have also shown how prior knowledge of gene function and variant effect predictions can be used to predict growth differences of different strains of *S. cerevisiae* (Jelier *et al*, 2011) and *E. coli* (Galardini *et al*, 2017). These analyses illustrate ways to calculate gene burden scores across different effect predictors. We found a significant but limited overlap between the gene-condition associations derived here with those found in gene KO studies in the reference lab strain. This small overlap could be due to a number of reasons including errors in variant effect predictions; limited sample size for the associations (i.e. 93 strains); epistatic interactions of variants and different protocols for fitness measurements. The effects of a genetic variation *in vivo* can be complex and depend on both genetic and environmental factors (Wray *et al*, 2013; Burga *et al*, 2011; Perez *et al*, 2017). Several studies have shown that many variants annotated as disease-causing or predicted as deleterious have been identified in healthy humans (Xue *et al*, 2012). In addition to these potential causes of error, it is assumed here that the loss of function of a given gene will have the same phenotypic consequence across individuals of the same species. The extent by which this assumption is true remains to be tested.

Despite the limitations discussed, given the growing number of efforts to sequence exome and genomes for panels of individuals, the incorporation of variant prioritization by different approaches into association analyses will become more prevalent. The mutfunc resource can provide such variant effect predictions with mechanistic annotations for 3 species. We illustrate how this resource can be applied in different scenarios and given the architecture used these analysis can be easily incorporated into large scale full genome or exome sequencing efforts.

## Methods

### Genetic variant data collection

A total of 896,772 genetic variants occurring in for 405 haploid and diploid *S. cerevisiae* strains were collected from four studies (Strope *et al*, 2015; Gallone *et al*, 2016; Bergström *et al*, 2014; Zhu *et al*, 2016). All but one study by Strope et al. provided processed variant calls in VCF format. Variants were called for the Strope et al. study using the following pipeline. Raw reads were obtained from the ENA resource (Leinonen *et al*, 2011). Adapter sequences were removed using cutadapt v1.8.1and reads were mapped to the *S. cerevisiae* genome version 64 using BWA-MEM v0.7.8 (https://arxiv.org/abs/1303.3997). Duplicate reads were discarded using picard v1.96 (https://github.com/broadinstitute/picard) and reads were realigned using the GATK indel realigner v3.3 (McKenna *et al*, 2010). Base alignment qualities were computed using samtools v1.2 (Li *et al*, 2009) and variants were called using freebayes v0.9.21-15-g8a06a0b and the following parameters `--no-complex, --genotype-qualities, --ploidy 1 and --theta 0.006`. The VCF was filtered for calls with `QUAL > 30, GQ > 30` and D`P > 4`. VCF for individual *S. cerevisiae* strains were combined and coding variants were called using the `predictCoding` function of the VariantAnnotation R package (Obenchain *et al*, 2014).

A total of 3,198,692 coding variants in *H. sapiens* for over 65,000 individuals was collected from the ExAC consortium along with corresponding adjusted allele frequencies. Ensembl transcript positions were mapped to UniProt by performing Needleman-Wunsch global alignment of translated Ensembl transcript sequences against the UniProt sequence using the `pairwiseAlignment` function in the Biostrings R package. The mapping between Ensembl transcript IDs (v81) and UniProt accessions was obtained from the biomaRt R package (Smedley *et al*, 2015). In the case that multiple alleles mapped to the sample single amino acid substitution, the one with the highest adjusted allele frequency was retained.

A total of 139,167 variants were obtained from ClinVar. Only variants that did not match one of the following clinical significance terms were removed: ‘Benign’, ‘Benign/Likely benign’, ‘Likely benign’, ‘Likely pathogenic’, ‘Pathogenic/Likely pathogenic’, and ‘Pathogenic’. Variants with a review status of ‘no assertion criteria provided’ were also removed, as those reflect variants that have been assigned clinical significance without any particular criteria. The final filtered set contained 39,597 variants. Of these variants, 44% were classified as pathogenic or likely pathogenic. For *S. cerevisiae*, a total of 8,083 manually curated variants were obtained from (Jelier *et al*, 2011), 34.5% (2,812) of which were labelled as deleterious. Variants were collected from a combination of the UniProt database (Apweiler *et al*, 2004), Protein Mutant Database (Kawabata *et al*, 1999), *Saccharomyces* genome database (Cherry *et al*, 2012) and mutations that are identified in essential genes (Liti *et al*, 2009).

### Essential genes

A total of 2,501 essential genes identified using gene trapping technology in two haploid *H. sapiens* cell lines KBM7 and HAP1 were obtained from (Blomen *et al*, 2015). These were further filtered for genes that were essential in both cell lines, for a total of 1,734 genes. A total of 1,156 essential genes in *S. cerevisiae* were obtained from the *Saccharomyces* Genome Deletion Project (Giaever *et al*, 2002).

### Predicting impact on protein stability and protein interaction interfaces

Experimentally determined structures were obtained from the protein data bank (PDB). Large structures that did not have a corresponding PDB file were downloaded in mmCIF format and converted to PDBs using the PyMOL Python library v1.2r3pre (pymol.org). Mapping of coordinates from PDB to UniProt residues was derived from the SIFTS database (Velankar *et al*, 2013). Structures with a resolution above 3 angstroms were discarded and a single representative structure maximising the coverage of the protein was retained. Homology modelling was carried out for proteins with no experimentally determined structures using ModPipe version 2.2.0 (Pieper *et al*, 2009) and the following parameters: `--hits_mode 1110 and --score_by_tsvmod OFF`. For each protein, the model with the highest normalised DOPE score was retained. Experimental and homology modelled structures for protein interactions were obtained from the Interactome3D database [23399932]. Relative solvent accessibility (RSA) for all residue atoms was computed using NACCESS for proteins individually, and in the interaction complex. Interface residues were defined as those with any change in RSA. All other calculation of RSA was carried out using freeSASA v1.1 (Mitternacht, 2016).

The impact of variant on stability was computed using FoldX v.4.0 (Schymkowitz *et al*, 2005). All structures were first split by chain into individual PDB files and repaired using the `RepairPDB` command, with default parameters. The `Pssm` command is then used to predict ΔG with `numberOfRuns=5`. This performs the mutation multiple times with variable rotamer configurations, to ensure the algorithm achieves convergence. The average ΔG of all runs is computed and the ΔΔG is computed as the difference between the wildtype and mutant. The impact of variants on interaction interfaces is measured similarly, with the exception of structures being provided in binary interaction, rather than individual chains.

### Predicting the impact of variants on PTMs and linear motifs

For *S. cerevisiae*, a total of 20,056 phosphosites and 2,219 kinase-substrate associations were obtained from the PhosphoGRID database (Sadowski *et al*, 2013). A total of 1,070 of other PTM sites was obtained from the dbPTM database (Lee *et al*, 2006). For *H. sapiens*, all PTM data, including that of phosphorylation and kinase-substrate associations were obtained from PhosphoSitePlus (Hornbeck *et al*, 2012), for a total of 296,147 sites. For *E. coli*, a total of 483 PTM sites were obtained from dbPTM (Lee *et al*, 2006). Linear motif data for *S. cerevisiae* and *H. sapiens*, including annotated linear motif binding sites and regular expression patterns, were obtained from the ELM database (Dinkel *et al*, 2016).

Impact of variants on phosphosites and flanking regions was measured using the MIMP algorithm (Wagih *et al*, 2015), with default parameters. For other PTMs, a variant was predicted to be impactful if it resulted in the change of the modified residue. For linear motifs, a variant was predicted to be impactful if it causes a loss of match for associated regular expression pattern.

### Predicting the functional impact of variants using conservation

All protein alignments were built against UniRef50 (Suzek *et al*, 2015), using the `seqs_chosen_via_median_info.csh` script in SIFT 5.1.1 (Ng & Henikoff, 2003). The siftr R package (https://github.com/omarwagih/siftr), an implementation of the SIFT algorithm, was used to generate SIFT scores with parameters `ic_thresh=3.25 and residue_thresh=2`.

### Transcription factor binding sites

A total of 177 *S. cerevisiae* TFs binding models were collected in form of a position frequency matrices (PFMs) from JASPAR (Sandelin *et al*, 2004) and converted to position weight matrices (PWMs) using the TFBSTools R package (Tan & Lenhard, 2016). PWMs were trimmed to eliminate consecutive stretches of low information content (<0.2) on either terminus. To identify genes likely regulated by a particular TF, a combination of TF-knockout expression and ChIP-chip experiments were used, as similarly described in (Gonçalves *et al*, 2017). Genome-wide gene expression profiles for 837 gene-knockout strains were obtained from three studies (Chua *et al*, 2006; Hu *et al*, 2007; Kemmeren *et al*, 2014), 148 of which were a known TF with a defined PWM. Studies provided either a Z-score or p-value for each gene as a measure of over or under-expression, relative to the distribution of values for all genes. Two-tailed p-values were computed from Z-scores when a p-value was not provided. In cases where TF knockout was repeated between studies, the lowest p-value for each gene was used. ChIP-chip tracks for 355 TFs were collected from four studies (Harbison *et al*, 2004; Rhee & Pugh, 2011; Tachibana *et al*, 2005; Venters *et al*, 2011) via the *Saccharomyces* genome database. Of the 355 of the TFs, 144 (56%) had a defined PWM. Potential binding sites were then only searched for in TF-gene pairs with a p-value below 0.01 and the corresponding ChIP-chip region upstream of the regulated gene. A normalised log score of 0.80 was used as the cutoff for defining putative binding sites. Similarly, for *H. sapiens*, 454 TF PWMs were generated from JASPAR PFMs. ENCODE clustered ChIP-seq data were obtained for 161 TFs, of which 72 had a PWM. Only those regions were scored against the corresponding PWM. For *E. coli*, a total of 1,905 TF-matching sequences across 84 TFs were obtained from RegulonDB (Gama-Castro *et al*, 2016) and used to construct PWMs. A total of 2,416 experimentally identified TFBS were obtained for 79/84 TFs from RegulonDB. These sites were used as putative binding sites for downstream variant predictions.

Potential target sequences were scored against the PWM using the log-scoring scheme defined in (Wasserman &Sandelin, 2004) and normalised to the best and worst matching sequence to the PWM. The resulting score lies between 0 and 1, where 1 signifies strong predicted binding by the factor, whereas 0 signifies predicted lack of binding. Potential binding sites were scored in the presence (S_wt_) and absence (S_mt_) of a variant. Three separate metrics are used to quantify the change in binding between the reference and alternate allele. The first one is simply the difference in the normalised log score, S_wt_ - S_mt_, where a large positive value indicates loss of binding. The second is the difference in binding percentile. Here, random oligonucleotides are used to generate a negative distribution of log normalised scores for each TF. The percentile of each wildtype p_wt_ and mutant scores p_mt_ is computed from this distribution, and the difference, p_wt_ - p_mt_, is used to quantify the magnitude of impact. The last is the difference in the relative information content. This can be thought of as the difference of letter height in a sequence logo. Given that the wildtype and mutant bases have relative frequencies of f_wt_ and f_mt_, respectively and a position has an IC value of γ, then this is computed as (f_wt_·γ) - (f_mt_·γ). This value ranges from 0 to 2, where 0 indicates little to no impact on a critical base, and 2 indicates a strong one.

### Implementation of mutfunc

Described predictors were used to precompute effects for all amino acid and nucleotide substitutions. The mutfunc web server at http://mutfunc.com uses the Java and Scala-based Play Framework v1.3.7 backend (http://www.playframework.org) along with a MySQL database. The front-end utilises a modified version of the the Twitter Bootstrap UI library (http://twitter.github.com/bootstrap). Visualization tools used include a modified version of the neXtProt feature viewer v0.1.52 (https://github.com/calipho-sib/feature-viewer) for interactive visualisation of protein sequence features, WebGL protein viewer v1.1 for interactive visualisation of protein structures v1.8.1 (https://github.com/biasmv/pv), and a modified version of the JSAV v.1.10 library (https://github.com/AndrewCRMartin/JSAV) for visualization of multiple sequence alignments.

### Chemical genetic screening

The screening was carried out in 1536 format on synthetic complete media with the addition of the appropriate chemical at a specific concentration. The Singer RoToR (Singer instruments, UK) was used to replicate screening plates in 1536 format. Agar plates were pinned onto the conditioned media and allowed to grow for 48 or 72 hours at 30 degrees centigrade (unless specified otherwise). Each experiment was replicated once for quality control. After incubation, plates were imaged and colony sizes were extracted using IRIS version v0.9.7 (Kritikos *et al*, 2017) with the “Colony growth” profile, which extracts colony size, circularity and opacity from each colony in each plate. Individual strains were scored using the E-MAP software, which transforms colony sizes into s-scores (Collins *et al*, 2006). In brief, a surface correction algorithm is applied to each plate, the outer frame effect is corrected by bringing the two outermost rows and columns to the plate middle median. All the plates are then normalized to the overall median, followed by a variance correction and finally the s-score calculation. The resulting s-scores are quantile normalized in each condition separately and final s-scores from both replicates are averaged.

### Calculating gene and complex disruption scores

Scores produced by different predictors were standardized in order to reflect the likelihood of identifying a deleterious mutation (P_del_). For SIFT, a curated gold standard set of 8,083 variants in 1,346 yeast genes with known tolerated or deleterious effects were obtained from Jelier et al. (Jelier *et al*, 2011). The negative natural logarithm of the SIFT score was binned by 0.5 and for each bin, the proportion of deleterious variants was computed. A binomial logistic regression was fit to the proportion values and used to compute subsequent Pdel values for subsequent SIFT scores. For FoldX, 964 gold-standard mutations across 34 experimentally identified proteins structures with both experimentally quantified ∆∆G values and FoldX-predicted ∆∆G values were obtained from Guerois et al. (Guerois *et al*, 2002). A variant was labelled destabilising if ∆∆G was greater than 1. Mutations were binned by predicted ∆∆G at intervals of 0.4 and for each bin, the proportion of destabilising variants was computed. A binomial logistic regression model was similarly fit to the data and used to compute subsequent P_del_ for FoldX-predicted ∆∆G values. For variants disrupting start or stop codons we assigned P_del_ value of 1. Since nonsense variants occurring closer to the C-terminal of a protein are less likely to impact function, we only assign P_del_ value of 1 for nonsense variants occurring in the first 50% of the protein, otherwise a value of 0 was used. Gene burden scores are then computed as the variant with the maximum P_del_ score and describes the predicted likelihood that a protein has an affected function (P_AF_). Similarly, for protein complexes the maximum P_del_ score for any complex subunit was selected to reflect the protein complex P_AF_ score. Variants with a MAF > 20% were considered unlikely to be deleterious given their high frequency in the population and were discarded prior to the burden score analysis.

### Genotype to phenotype association analysis

The associations were carried out using the MatrixEQTL R package (Shabalin, 2012) with the modelLINEAR mode. The significance of the association was measured using a t-statistic. For the associations genes and complex binarized P_AF_ scores were used as genotypes where a P_AF_ score above or below 0.9 is given a value 1 and 0, respectively and growth phenotypes are used in lieu of gene expression. A *p-value* threshold of 0.001 was used for all associations and multiple testing correction was carried out using the false discovery method. Effect size was computed using Glass’ ∆. For the case (p) and control (n) group, differences in the mean was computed relative to the standard deviation of one of the groups. Given the mean (μ_i_) and standard deviation (σ_i_) for a given group i this is computed as ∆i = (μ_p_ − μ_n_)/σ_i_. For robustness this was computed in both direction and the final effect size, ∆, is reported as the minimum absolute value of effect sizes in both directions.

---

**Supplementary figure 1.**
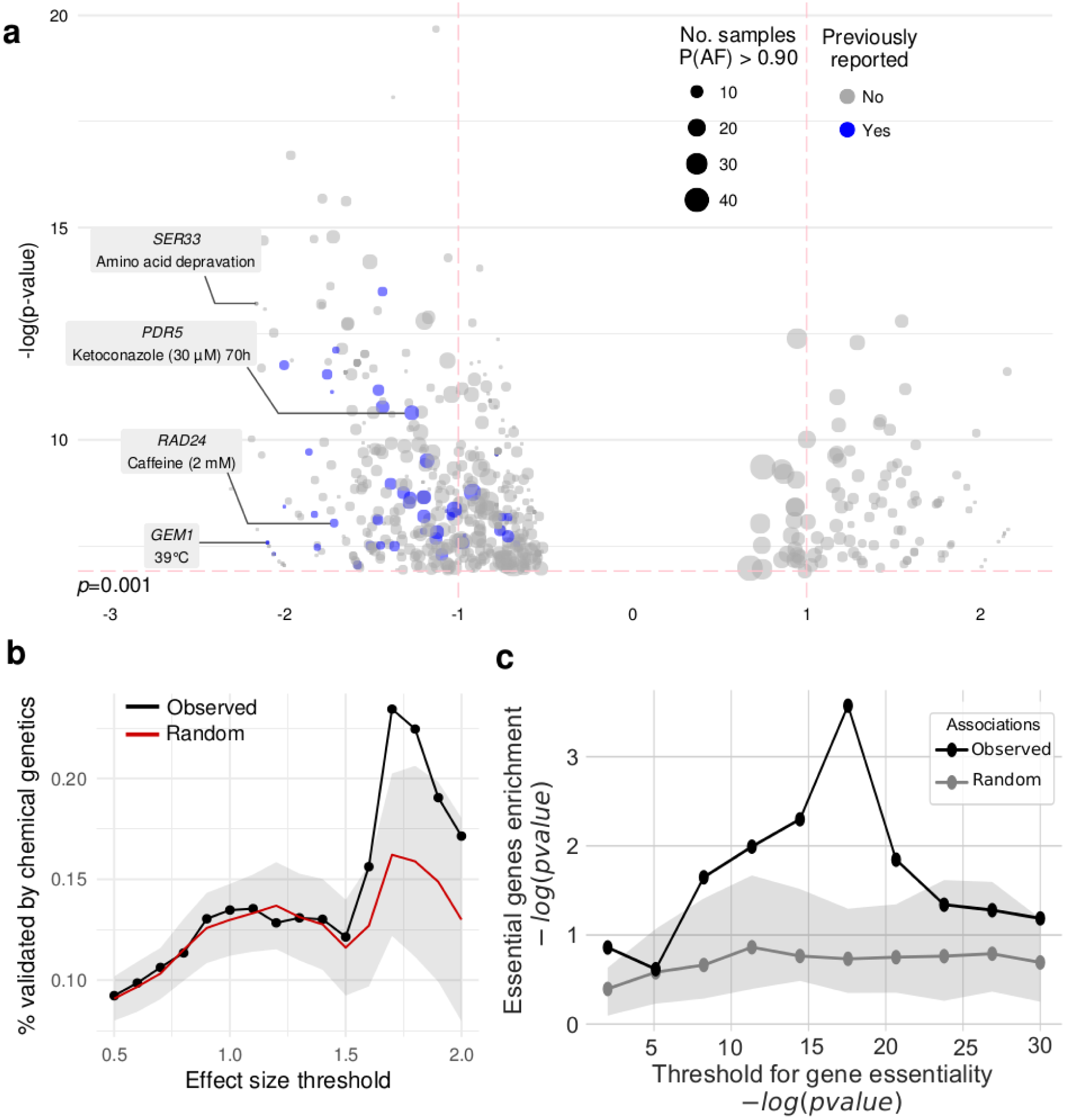
Gene and protein complex level phenotype association analysis show significant but modest enrichment in prior knowledge of gene KO growth phenotypes. (a) Gene to phenotype associations using the gene burden scores are preferentially negative. (b) the fraction of gene-phenotype associations that is validated by chemical genetic information derived from gene deletion experiments. The significance of the observed overlap was tested using permutation testing and only found to be significant at effect size threshold. (c) Associations between protein complexes and conditions were benchmarked by calculating the enrichment of previously known gene-condition associations from gene deletion studies. An enrichment was observed for some cut-offs for the gene-deletion condition dependent essentiality but only found to be better than random expectation for stringent cut-off.

**Supplementary table 1 - Growth measurements for *S. cerevisiae* strain panel.** The growth measurements expressed as s-scores for each strain in each condition is listed.

